# Trem2^R47H^ and reduced *TREM2* expression both mimic human Alzheimer’s disease signatures in mice

**DOI:** 10.64898/2025.12.17.694758

**Authors:** Tamar R. Abel, Ravi S. Pandey, Annat Haber, Dylan Garceau, Michael Sasner, Kevin P. Kotredes, Gregory A. Cary, Adrian Oblak, Gareth R. Howell, Bruce T. Lamb, Gregory W. Carter

**Author notes:** **Correspondence:** Gregory W. Carter, The Jackson Laboratory, 600 Main Street, Bar Harbor, Maine, 04609 USA.

## Abstract

**INTRODUCTION:** *TREM2* loss of function variants are associated with late-onset Alzheimer’s disease (LOAD). We molecularly assessed mice with the missense variant, R47H (Trem2*R47H^HSS^), and mice with additional cryptic splicing and reduced *Trem2* expression (Trem2*R47H) while comparing relevance to human LOAD.

**METHODS:** The aberrant splice acceptor site in the Trem2*R47H mouse was humanized resulting in the Trem2*R47H humanized splice site (Trem2*R47H^HSS^) mouse. RNA sequencing was performed on mouse brain tissue and signatures were compared to human postmortem brain expression in LOAD cohorts.

**RESULTS:** Trem2*R47H mice had alternative splicing leading to reduced *Trem2* expression; Trem2*R47H^HSS^ mice expressed *Trem2* at wild-type transcript and protein levels. Both models correlated with similar LOAD-associated signatures, and had similar effects on immune response, synapse, and vasculature biodomains. The Trem2*R47H model additionally affected extracellular matrix and myelination signatures.

**DISCUSSION:** We demonstrated that Trem2*R47H and Trem2*R47H^HSS^ mice are complementary models for the study of molecular contributions to LOAD pathology.

## 1 Introduction

Triggering receptor expressed on myeloid cells 2 (*TREM2*) is a risk gene for Alzheimer’s disease (AD) [1, 2] and other neurodegenerative diseases [3–6]. *TREM2* is highly expressed by microglia in the brain where it binds to lipid ligands, promotes phagocytosis, and influences the release of cytokines that modulate inflammation [7, 8]. Microglia are involved in processes of amyloid plaque response, synaptic pruning, activation of glial cell populations, and myelination indicating that *TREM2* may play a causal role in AD pathogenesis [9, 10]. However, the mechanisms by which *TREM2* influences neurodegenerative disease are poorly understood.

The *TREM2*\*R47H variant, a rare missense mutation in exon 2, is one of the strongest genetic risk factors associated with late onset AD (LOAD) [1, 2]. Cellular and structural studies have indicated a partial loss of function due to R47H which leads to impaired ligand binding, reduced activation, and less downstream kinase signaling [11, 12]. In contrast, a recent study identified a gain of function variant which increases *TREM2* expression and is associated with reduced AD disease risk [13]. Another study demonstrated that many rare recessive missense *TREM2* variants associated with AD lead to alternative splicing including exon skipping, suggesting that *TREM2* mis-splicing may contribute to AD pathology as well [14]. In context of *Trem2* knock-out (KO) studies in murine models [15, 16] these findings suggest that *TREM2* loss of function by any mechanism is likely to influence AD risk. However, it is important to note that not all studies associate *TREM2* loss of function with increased AD pathology [17], and individual impact on LOAD may depend on multiple factors including disease stage and genetic background.

There are several murine models for Trem2*R47H [18–20]. The Model Organism Development and Evaluation for Late-onset Alzheimer’s Disease (MODEL-AD) center at the Indiana University, The Jackson Laboratory, and the University of Pittsburgh (IU/JAX/PITT) has successfully created and characterized one of these models [20–22]. Early findings indicated that Trem2*R47H mice had a phenotype consistent with that of *Trem2* haploinsufficiency [18]. Multiple groups subsequently identified a cryptic splice site that results in the introduction of a premature stop codon and expression of a novel murine transcript with a truncation at the 5’ end of exon 2 [19, 21, 23] – ultimately resulting in decreased *Trem2* expression [19, 20]. While this complicates the study of AD-risk associated with the R47H missense variant (human *TREM2* R47H does not alter expression [19]), it also provides an alternative model for studying the relevance of *TREM2* expression levels and alternative splicing in AD-associated pathological molecular signatures.

There are few published models which represent the Trem2*R47H missense variant without any loss of gene expression. One includes a Rat *Trem2*\*R47H model [24] which was reported to have *Trem2* expression and protein levels consistent with the wild-type rat. Additionally, MODEL-AD center at the University of California Irvine created the Trem2*R47H Normal Splice Site [2] mouse by introducing synonymous mutations to correct the cryptic splice site. These mice lack the altered splice site, novel transcript isoform, and reduced expression present in Trem2*R47H mice [23].

Our goal was to molecularly compare a new humanized model of Trem2*R47H (with no loss of *Trem2* expression) to the existing Trem2*R47H IU/JAX/PITT MODEL-AD model. Given the literature, we hypothesized that both *Trem2* models would demonstrate molecular associations with LOAD signatures. Beginning with Trem2*R47H mice we reverted the synonymous mutations associated with the cryptic splice site by humanizing the exon 2 splice site boundary – dubbed the Trem2*R47H humanized splice site (Trem2*R47H^HSS^) mouse. Using bulk transcriptomic data from B6 controls, Trem2*R47H mice, and our new Trem2*R47H^HSS^ model we validated *Trem2* expression levels. Next, we molecularly characterized each model and determined LOAD-associated gene expression signatures. While both *Trem2* models had overlap, the Trem2*R47H mice demonstrated additional correlation with LOAD-associated extracellular matrix and myelination signatures. Overall, we determined LOAD signatures for two Trem2*R47H models and demonstrated that both models are suitable for the study of *Trem2*-associated contributions to LOAD pathology.

## 2 Methods

### 2.1 Development of Trem2 mouse models

Two *Trem2* mouse alleles were generated using direct delivery of CRISPR-Cas9 reagents to mouse zygotes. Analysis of genomic DNA sequence surrounding the target region using the Benchling guide RNA design tool identified guide RNA sequences with a suitable target endonuclease site near the region of interest. *Streptococcus pyogenes* Cas9 (SpCas9) V3 protein and gRNAs were purchased as part of the Alt-R CRISPR-Cas9 system using the crRNA:tracrRNA duplex format as the gRNA species (IDT, USA). Alt-R CRISPR-Cas9 crRNAs (Product# 1072532, IDT, USA) were synthesized using the gRNA sequences specified in the below section and hybridized with the Alt-R tracrRNA (Product# 1072534, IDT, USA) as per manufacturer’s instructions. These guide RNAs were evaluated for their ability to induce double stranded DNA cleavage events in a mouse kidney cell line.

The sequence of the single stranded DNA Oligonucleotide (ssDO),that functions as the DNA repair template, was designed based on the SpCas9 cleavage and repair model described by Corn and colleagues [25]. Features included asymmetric homology arm lengths, flanking the desired SNPs, as well as being the antisense sequence in reference to the gRNA. Repair oligonucleotides were synthesized by IDT (USA).

The R47H mutation was introduced into the mouse *Trem2* locus in the C57BL/6J (JAX #664) background by cutting with a CRISPR guide (5’-GAAGCACTGGGGGAGACGCA-3’) just upstream of the area of interest. The R47H allele was introduced using a repair oligo (183 nucleotides) containing a nucleotide G>A point mutation (at nucleotide 89 in this oligo sequence) for the amino acid sequence change at R47H and 2 silent mutations (lysine AAG>AAA (at nucleotide 93 in this oligo sequence) and alanine GCC>GCA (at nucleotide 96 in this oligo sequence)) into the gene to prevent re-cutting of the donor sequence in homologous directed repair.

Sequence of repair oligonucleotide: GCCCTCAACACCACGGTGCTGCAGGGCATGGCCGGCCAGTCCTTGAGGGTGTCATGTACT TATGACGCCTTGAAGCACTGGGGGAGACACA*AaGCaT*GGTGTCGGCAGCTGGGTGAGGAG GGCCCATGCCAGCGTGTGGTGAGCACACACGGTGTGTGGCTGCTGGCCTTCCTGAAGAA GCGG

This model is available from the JAX mouse repository and is referred to as “Trem2*R47H” or C57BL/6J-Trem2^em1Adiuj^/J (JAX #27918).

The R47H^HSS^ model was created by using CRISPR to edit sites in the existing Trem2*R47H model. Unique sites in this model were targeted using the CRISPR guide 5’-GGAGACACAAAGCATGGTGT-3’, and sequences to humanize the cryptic splice site just 3’ of the R47H mutation were incorporated with the repair oligonucleotide:

CCCATTCCGCTTCTTCAGGAAGGCCAGCAGCCACACACCGTGTGTGCTCACCACACGCTGGCATGGGCCCTtCTCtCCCAGCTGgCGgCACCAgGCcTTGtGTCTCCCCCAGTGCTTCAAGGCGTCATAAGTACA.

This model is available from the JAX mouse repository and is referred to as “Trem2*R47H^HSS^” or B6.Cg-Trem2^em4Adiuj^/J (JAX #33781).

### 2.2 Animal housing

C57BL/6J (B6), B6.*Trem2*R47H*, and B6.*Trem2*R47H^HSS^*animals were bred, room-controlled, and aged in the main Research Animal Facility (RAF) located in Center for Biometric Analysis (CBA) building. The dedicated housing consists of PIV caging, wood shavings bedding (aspen or pine), enrichment (nestlets and huts), with temperature controlled at a setting of 72±2°F and humidity at 50±20%. The facility was on a 12:12 L:D schedule (lights on at 6:00 am). All mice were fed LabDiet® 5K52/5K67 (contains 6% fat) diet and water ad lib.

### 2.3 Genotyping

Founder mice with the appropriate sequence were identified using a nested PCR approach, with initial product spanning the region upstream and downstream of the repair construct, and internal primers covering the sequence of the repair construct. PCR products were sequenced to verify sequences. Founder mice were bred to C57BL/6J mice (JAX #664) for at least two generations, then heterozygotes were intercrossed to create homozygote cohorts for analysis.

For the Trem2*R47H model, primers used were:

Forward-1: TTC CAA GCA AGT GGC TGT CT

Reverse-1: TTC CAG CAA GGG TGT CAT C

For the Trem2*R47H^HSS^ model, primers used were:

Forward-1: CTG CAC TCC TGG AAC TGC TT

Forward-2: CTC GGA GAC TCT GAC ACT GG

Once colonies were established, end point analysis assays were used to genotype litters.

For the Trem2*R47H model, oligonucleotides used were:

Forward primer: ACT TAT GAC GCC TTG AAG CA

Reverse primer: CTT CTT CAG GAA GGC CAG CA

Wild-type probe: AGA CGC AAG GCC TGG TG

Mutant probe: AGA CAC AAA GCA TGG TGT CG

For the Trem2*R47H^HSS^ model, oligonucleotides used were:

Forward primer: CTT GAA GCA CTG GGG GAG AC

Reverse primer: TCT TCA GGA AGG CCA GCA G

Wild-type probe: TGT CGG CAG CTG GGT GA

Mutant probe: TGC CGC CAG CTG GGA G

For detailed PCR protocols, see JaxMice datasheets:

Trem2*R47H: https://www.jax.org/strain/027918

Trem2*R47H^HSS^: https://www.jax.org/strain/033781

### 2.4 Animal anesthesia, perfusion, and brain tissue collection

Upon arrival at the terminal endpoint for each aged mouse cohort, individual animals were weighed prior to intraperitoneal administration of either: (A) ketamine (100mg/kg) and xylazine (10mg/kg); or (B) tribromoethanol (1mg/kg). Routine confirmation of deep anesthesia was performed every 5 minutes by toe pinch. Animals were perfused transcardially with 1xPBS solution to clear the vascular system of all blood (left ventricle to right atrium with approximately 10mL of 1xPBS). Completion of perfusion and clearance of the vascular system was indicated by a blanching of the liver. Anesthetized and subsequently perfused animals were decapitated, and skulls submerged in cold 1xPBS. The whole brain was carefully removed from the skull, weighed, and divided midsagitally, into left and right hemispheres, using a brain matrix. Complete brain hemispheres were quickly aliquoted into cryotubes and immediately snap frozen on liquid nitrogen, and stored long-term at -80°C.

### 2.5 Western blot

Mice were euthanized, and hemibrains were rapidly frozen on ice. Samples were homogenized in RIPA buffer supplemented with protease and phosphatase inhibitors (ThermoFisher Halt™). Lysates were incubated on ice for 30 minutes and centrifuged at 14,000 × g for 15 minutes at 4°C. Supernatants were collected, and total protein concentration was measured using the BCA Protein Assay Kit (ThermoFisher Scientific). Equal amounts of protein (20 µg per lane) were denatured in Laemmli buffer with β-mercaptoethanol and separated by SDS-PAGE on 10–12% polyacrylamide gels. Proteins were transferred to PVDF membranes (Millipore) using a semi-dry transfer system. Membranes were blocked in 5% non-fat dry milk in TBS-T (Tris-buffered saline with 0.1% Tween-20) for 1 hour at room temperature and incubated overnight at 4°C with primary antibodies against the target protein(s) of interest (anti-TREM2, anti-β-actin for loading control). After washing, membranes were incubated with HRP-conjugated secondary antibodies for 1 hour at room temperature. Signal was detected using enhanced chemiluminescence (ECL, GE Healthcare) and visualized with a ChemiDoc™ Imaging System (Bio-Rad). Band intensities were quantified using ImageJ software (NIH). Target protein expression levels were normalized to β-actin. Data were analyzed using one-way ANOVA followed by Tukey’s post hoc test for multiple comparisons. A p-value < 0.05 was considered statistically significant.

### 2.6 RNA Isolation, preparation, and sequencing

Total RNA was isolated from tissue using the NucleoMag RNA Kit (Macherey-Nagel) and the KingFisher Flex purification system (ThermoFisher). Tissues were homogenized in MR1 buffer (Macherey-Nagel) using a Bead Ruptor Elite (Omni International). RNA isolation was performed according to the manufacturer’s protocol. RNA concentration and quality were assessed using the Nanodrop 8000 spectrophotometer (Thermo Scientific) and the RNA ScreenTape Assay (Agilent Technologies).

Stranded libraries were constructed using the KAPA mRNA HyperPrep Kit (Roche Sequencing and Life Science), according to the manufacturer’s protocol. Briefly, the protocol entails isolation of polyA containing mRNA using oligo-dT magnetic beads, RNA fragmentation, first and second strand cDNA synthesis, ligation of Illumina-specific adapters containing a unique barcode sequence for each library, and PCR amplification. The quality and concentration of the libraries were assessed using the D5000 ScreenTape (Agilent Technologies) and Qubit dsDNA HS Assay (ThermoFisher), respectively, according to the manufacturers’ instructions. Libraries were sequenced 150 bp paired-end on an Illumina NovaSeq X Plus using the 10B Reagent Kit.

### 2.7 Bioinformatic analysis

All bioinformatic analyses were carried out using R (v4.4.1). All plots created using ggplot2 (v3.5.1) unless otherwise noted. Individual analyses are described in more detail in the following subsections.

#### 2.7.1 RNA-seq data preprocessing and quality control

RNA-Seq data were processed using nf-core/rnaseq pipeline [26]. Briefly, the sequence quality of reads was assessed using FastQC. Low-quality bases were trimmed from sequencing reads using Trim Galore. Reads were then aligned to the reference mouse genome (version GRCm38.p6) using STAR [27], and gene expression was quantified with RSEM [28]. RNA-seq data was used to confirm both the reported genotype and the sex of the mice. Mice with inconclusive genotype were excluded from the downstream analysis. A custom isoform was used to align the *Trem2* novel transcript as previously described [21].

#### 2.7.2 Trem2 transcript isoform and effect analysis

We plotted expression of *Trem2* transcript isoforms as either log transcripts per million (TPM) (all *Trem2* transcripts) or TPM (for individual transcripts due to low expression and presence of zero values) and p-values were calculated for each comparison using a Welch two sample t-test. Significance threshold for raw p-values was set at 0.05.

#### 2.7.3 Differential gene expression analysis

All differential expression analysis was carried out using DESeq2 (v1.44.0). Initial analysis was performed using age and sex stratified data with a linear regression analysis to regress out batch by including as a covariate. Additional analysis was carried out in twelve-month-old animals controlling for sex by including as a covariate in the linear regression model. Analyses were conducted separately using either the B6 gene expression as the reference or Trem2*R47H^HSS^ gene expression as the reference. Significance threshold for Benjamini–Hochberg adjusted p-values was set at alpha < 0.05. Histograms of p-value distribution were obtained from raw p-values.

#### 2.7.4 Trem2 model RNAseq correlation analysis

Pearson correlation coefficient was calculated for all transcripts (unfiltered for significance) from sex-matched differential expression of twelve-month-old Trem2*R47H and Trem2*R47H^HSS^ mice compared to B6 mice.

#### 2.7.5 Trans-species correlation analysis with human AD modules

Trans-species correlation analysis was carried out as previously described [29, 30]. Briefly, mouse fold-change was obtained from the regression coefficients calculated in the differential gene expression analysis step for three different comparisons: Trem2*R47H vs. B6, Trem2*R47H^HSS^ vs. B6, and Trem2*R47H vs. Trem2*R47H^HSS^. Pre-defined gene modules and human AD-specific fold-changes were obtained from the Accelerating Medicines Partnership Program for Alzheimer’s Disease (AMP-AD) modules [31]. These previously identified AMP-AD transcriptomic modules are enriched in AD as determined through harmonized co-expression or co-abundance analysis of omic data derived from human brain tissue and have been annotated for cellular function. Fold-change for each of the human AD-enriched genes in these modules was calculated previously comparing AD patient gene expression relative to healthy control gene expression. Using these gene expression data from both mouse and human we then determined the Pearson correlation between the effects of AD (vs. controls) in humans and the effects of the mouse perturbation (vs. wild type) in mice for each individual module [29]. All genes were unfiltered for significance. Significance of the correlation was defined as p-value < 0.05. Age and sex stratified mice were compared to sex-specific gene expression changes in the AMP-AD modules (i.e. fold-change for genes in the AMP-AD modules was calculated using either male- or female-only human populations).

#### 2.7.6 Subtype correlation analysis with human AD subtypes

Subtype correlation analysis also leveraged Pearson correlation between the effects of AD (vs. controls) in humans and the effects of the mouse perturbation (vs. wild type) in mice. Mice perturbations were once again obtained from the regression coefficients calculated in the differential gene expression analysis for three different comparisons: Trem2*R47H vs. B6, Trem2*R47H^HSS^ vs. B6, and Trem2*R47H vs. Trem2*R47H^HSS^. However, human differential expression data of AD patients vs. healthy controls was obtained from gene expression sets associated with pre-determined AD molecular subtypes reported in *Milind et al.* and *Neff et al.* publications [32, 33]. All genes were unfiltered for significance.

#### 2.7.7 Biodomain and sub-biodomain analysis

The correlation between effects of AD (vs. controls) in humans and the effects of the different gene variants in mice was also computed on sub-sets of genes annotated to different biological domains [34]. The biological domains (biodomains) represent different molecular endophenotypes consistently associated with human disease and genes are annotated to each domain via sets of largely non-overlapping gene ontology terms that are annotated to each domain. Each domain has also been clustered by disease signature to yield smaller groups of terms (subdomains) that are more specific sub-processes within each biodomain (Wiley et al, in prep). For each functional grouping defined by biodomains and subdomains, Pearson correlation between human log fold-change values computed from a meta-analysis of all AMP-AD transcriptome data [34] and mouse effects obtained from the regression coefficients calculated in the differential expression analysis for the three comparisons: Trem2*R47H vs. B6, Trem2*R47H^HSS^ vs. B6, and Trem2*R47H vs. Trem2*R47H^HSS^.

## 3 Results

### 3.1 Trem2*R47H^HSS^ mice maintain wild-type levels of *Trem2* expression with no novel isoform expression

Trem2*R47H mice contain synonymous mutations near the site of the engineered R47H mutation (**Figure 1A)** resulting in a cryptic splice site, introduction of a premature stop codon, and production of a novel truncated transcript which ultimately leads to decreased expression of the full *Trem2* transcript **(Figure 1B)**. For this study, we used the existing Trem2*R47H mouse [20] and also created a mouse model containing the R47H mutation in which the sequence surrounding the splice site was humanized (**Figure 1A**). This resulted in humanized splice site mice (Trem2*R47H^HSS^) which lack the synonymous mutations and cryptic splice acceptor site in exon 2.

**Figure 1.**
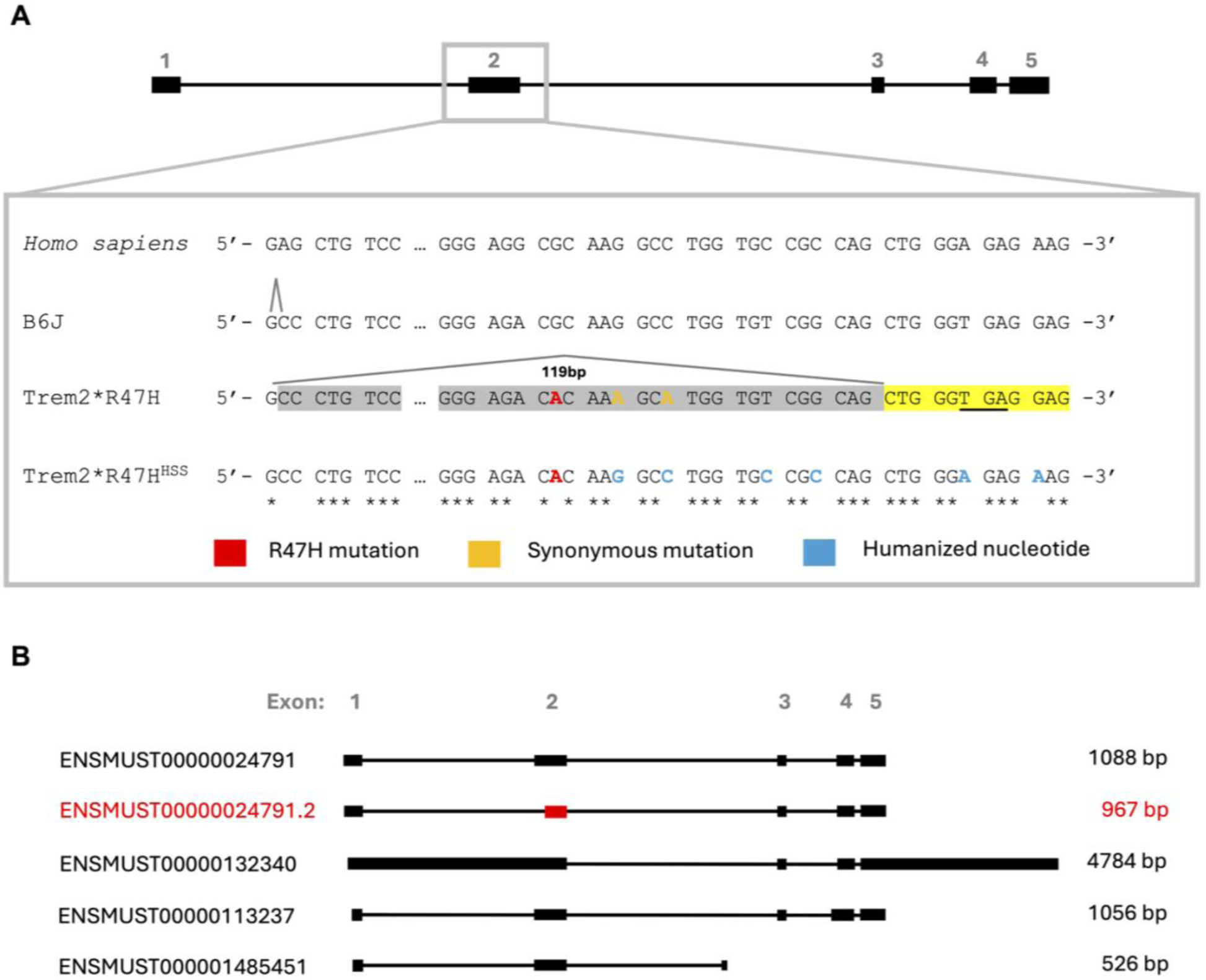
Diagrammatic representation of Trem2 mouse models and Trem2 transcripts. A) Aligned 5’ sequence of Trem2 exon 2. The human sequence is included for reference. Trem2*R47H mice contain knock-in of the R47H Alzheimer’s risk allele (red), synonymous mutations (yellow), and alternative splice site (yellow highlight). The premature stop codon (underlined) results in a truncated transcript with loss of 119bp region (grey highlight) and nonsense mediated decay of Trem2 transcript. The Trem2*R47H^HSS^ mice have a humanized splice site (blue nucleotides) that removes the synonymous mutations present in the Trem2*R47H mouse, thus resulting in normal Trem2 splicing and expression. B) Trem2 transcripts identified in Trem2*R47H mice including the primary transcript (ENSMUST00000024791) and the novel truncated isoform (ENSMUST00000024791.2) associated with a cryptic splice site in exon 2 introduced through synonymous mutations.

We created a cohort containing B6 controls, Trem2*R47H mice, and Trem2*R47H^HSS^ mice that were processed for bulk RNA-seq (**Table 1**). Both male and female as well as homozygous and heterozygous Trem2 mice were also included in the cohort, but heterozygous mice were only used in supplemental figures for comparison with homozygous Trem2*R47H and Trem2*R47H^HSS^ mice to determine if there was a dosing effect. As previously reported [20], Trem2*R47H mice had decreased expression of *Trem2* (**Figure 2A**) with significantly reduced expression of the primary transcript (**Figure 2B**), and concurrently an approximately three-fold increased expression of the novel truncated murine transcript as compared to B6 mice (**Figure 2C).** All other *Trem2* transcripts lacked any significant fold-change in Trem2*R47H mice. Of note, significantly decreased expression of *Trem2* was only present in homozygous, but not heterozygous, Trem2*R47H mice (Supplementary Figure 1-2). While heterozygous mice trended toward higher expression of the novel isoform the results were not statistically significant.

**Table 1.**
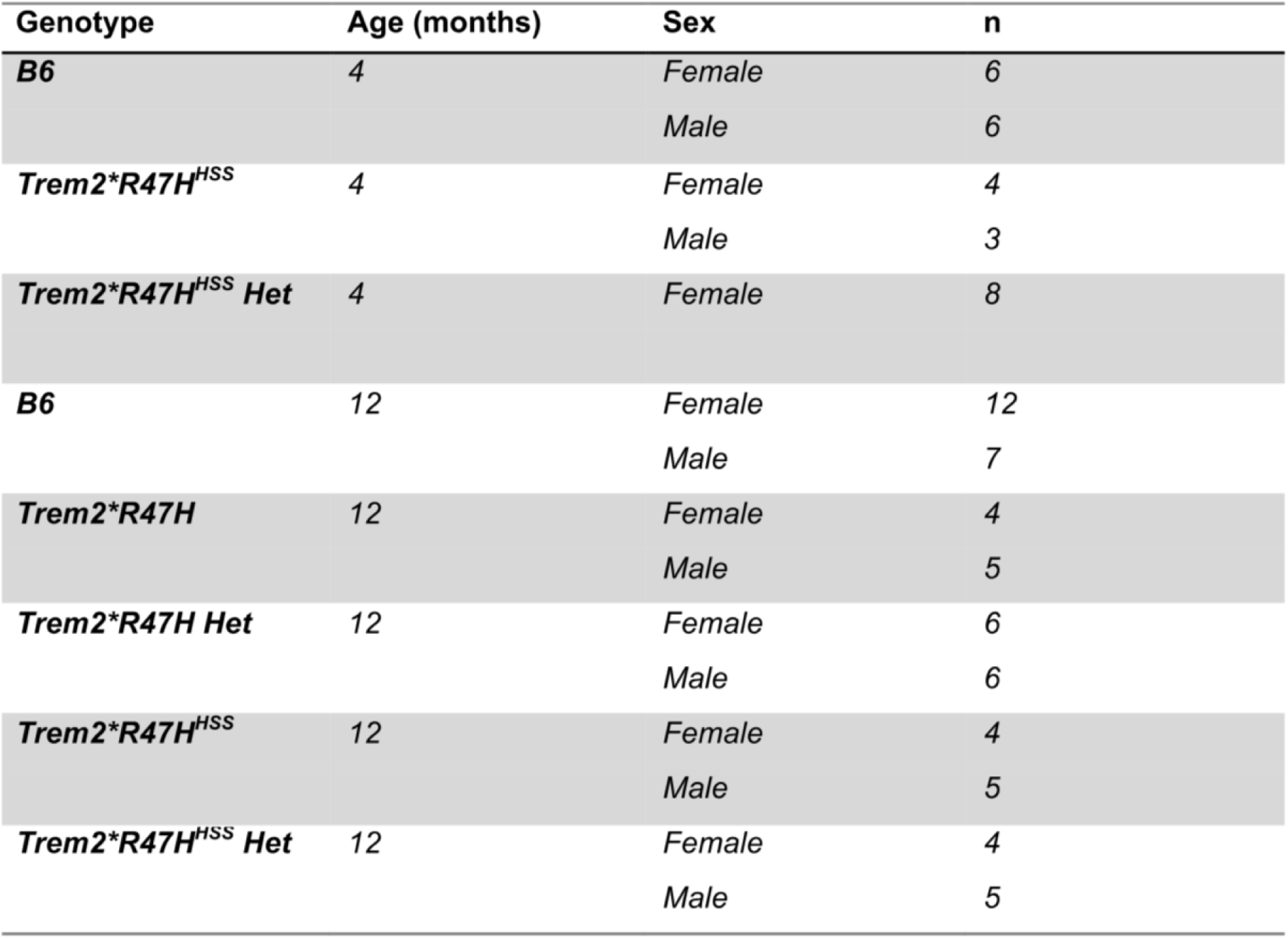
Trem2*R47H and Trem2*R47H^HSS^ mouse cohort.

| **Genotype** | **Age (months)** | **Sex** | **n** |
| --- | --- | --- | --- |
| ***B6*** | **4** | *Female* | 6 |
|  |  | *Male* | 6 |
| ***Trem2\**R47H*<sup>HSS</sup>*** | **4** | *Female* | 4 |
|  |  | *Male* | 3 |
| ***Trem2\**R47H*<sup>HSS</sup> Het*** | **4** | *Female* | 8 |
| ***B6*** | **12** | *Female* | 12 |
|  |  | *Male* | 7 |
| ***Trem2\**R47H**** | **12** | *Female* | 4 |
|  |  | *Male* | 5 |
| ***Trem2\**R47H* Het*** | **12** | *Female* | 6 |
|  |  | *Male* | 6 |
| ***Trem2\**R47H*<sup>HSS</sup>*** | **12** | *Female* | 4 |
|  |  | *Male* | 5 |
| ***Trem2\**R47H*<sup>HSS</sup> Het*** | **12** | *Female* | 4 |
|  |  | *Male* | 5 |

**Figure 2.**
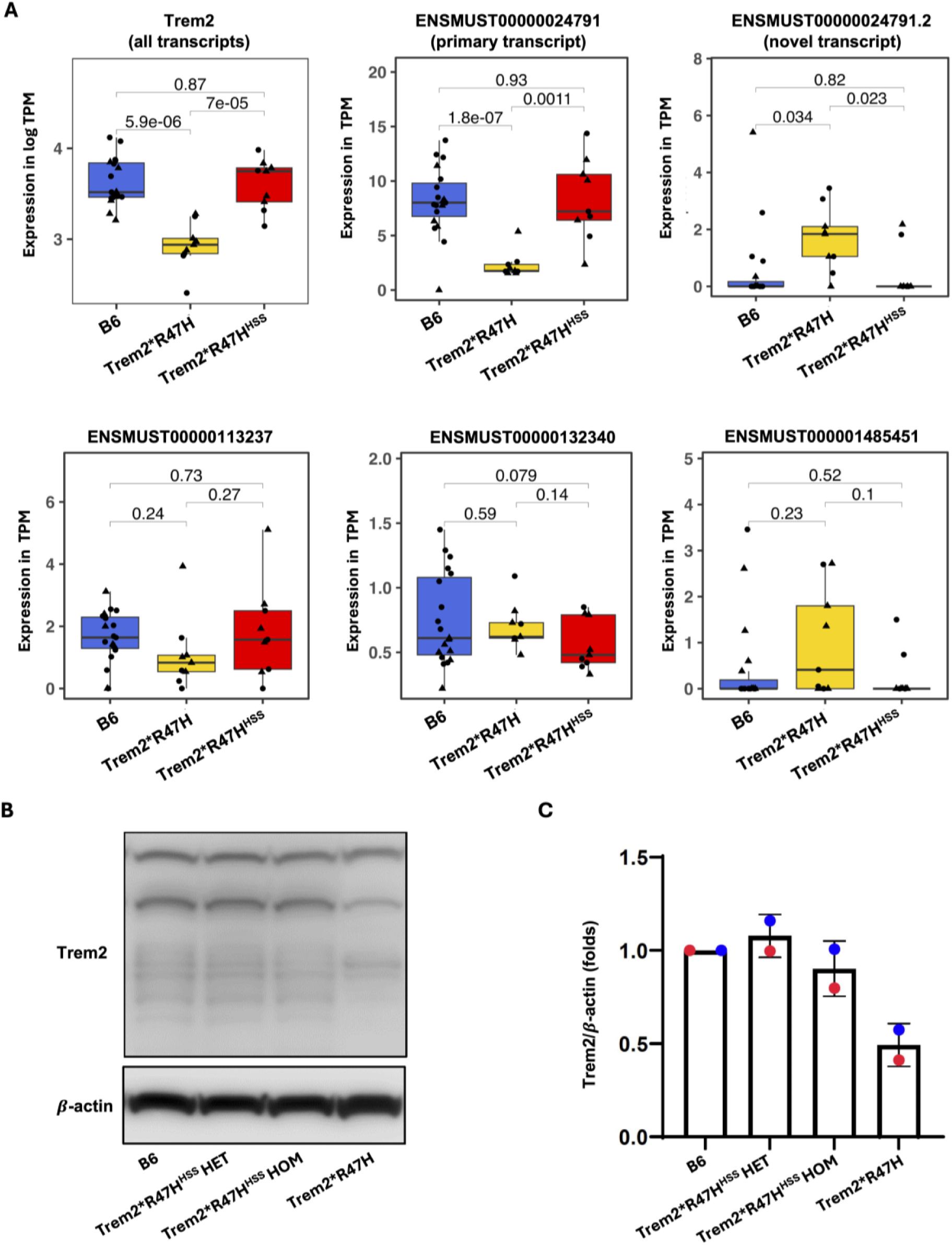
Trem2*R47H mouse with humanized splice site has normal Trem2 expression levels and does not express novel murine transcript. A) Boxplots showing the transcriptomic expression of Trem2 isoforms in B6 control mice (blue) as well as Trem2*R47H (yellow) and Trem2*R47H^HSS^ (red) mice. The sex is indicated by the shape of the dot (male = triangle, female = round). P-value for comparisons is included above boxplots. B) Western blot of Trem2 protein for B6, Trem2*R47H^HSS^ heterozygous (HET) and homozygous (HOM), and Trem2*R47H mice (all male). Beta-actin serves as a control for normalization. C) semi-quantitative analysis of band density (blue = male, red = female). Trem2*R47H mice showed 50% reduction in protein while Trem2*R47H^HSS^ maintained B6 levels of Trem2 expression.

In contrast, the Trem2*R47H^HSS^ mouse showed overall *Trem2* expression comparable to that of the B6 mice (**Figure 2A**) with highest expression of the primary transcript (**Figure 2B**). Neither Trem2*R47H^HSS^ or B6 mice expressed significant levels of the novel murine *Trem2* transcript (**Figure 2C**). Western blot results confirmed that Trem2 protein was present at wild-type levels in Trem2*R47H^HSS^ mice, and comparable to that of B6 mice (**Figure 2C-D**). Effect analysis on *Trem2* expression levels and *Trem2* transcript isoforms confirmed that regardless of sex, age, or genotype of the mice, the Trem2*R47H genotype was the only factor that resulted in a significant decrease of *Trem2* transcripts (in particular, the primary transcript) and increase of the novel *Trem2* isoform (Supplementary Figure 3).

### 3.2 Trem2*R47H and Trem2*R47H^HSS^ mice have similar transcriptomic profiles

To better understand differences on a systems level between Trem2*R47H and Trem2*R47H^HSS^ mice at twelve months, we performed differential expression analyses with either B6 mice as the reference or direct comparison of Trem2*R47H to Trem2*R47H^HSS^. Consistent with previous findings, *Trem2* showed neither a significant adjusted p-value or notable fold change for any comparisons between Trem2*R47H^HSS^ and B6 mice; and while significance of *Trem2* reduced expression in Trem2*R47H mice did not survive population stratification by sex or age of animals the Log2 fold-change was consistent with previous findings (Trem2*R47H vs. B6 male = -0.63, Trem2*R47H vs. B6 female = -0.40). Also, in keeping with these trends, the largest number of significantly differentially expressed genes were identified in comparisons between *Trem2* mouse models and B6 reference in analyses stratified by age and sex (**Table 2**). Notably, there were a greater number of significant genes identified in comparisons with male mice than female mice. There were very few significant genes associated with the direct comparison of Trem2*R47H and Trem2*R47H^HSS^ mice. Significant upregulated genes in Trem2*R47H male mice were *Ldlr* (p = 0.024, log2 fold-change (FC) = 0.40), Hgmcs1 (p = 0.024, log2 FC = 0.39), and Col3a1 (p = 0.041, log2 FC = 1.07) while the only significantly downregulated gene was a pseudogene. All genes identified in female mice were pseudogenes, apart from the significant upregulation of the mitochondrial gene Nd4l (p = 0.0079, log2 FC = 0.79). There was no overlapping differentially expressed genes between any of the mouse models. When observing the raw p-values we note that the distribution is relatively even, suggesting that there is an absence of statistically significant differences at the individual gene level between these two *Trem2* models (Supplemental Figure 4). However, when we performed a simple correlation of all genes in Trem2*R47H or Trem2*R47H^HSS^ mice as compared to the B6 reference (not filtered for significance), Trem2*R47H and Trem2*R47H^HSS^ showed significant correlations across all transcripts for both male and female mice (**Figure 3**). In this analysis of combined males and females *Trem2* was the only significant differentially expressed gene (identified in sex-corrected linear regression analysis of Trem2*R47H vs. B6 gene expression).

**Table 2.** Differentially expressed genes in Trem2*R47H and Trem2*R47HHss mice stratified by sex.

| Age | Comparison | Sex | Up DEGs | Down DEGs |
| --- | --- | --- | --- | --- |
| **4 mo** | *Trem2* * *R47H* <sup>HSS</sup><br>vs. B6 | Male | 0 | 4 |
|  |  | Female | 1 | 2 |
| **12 mo** | *Trem2* * *R47H* <sup>HSS</sup><br>vs. B6 | Male | 127 | 175 |
|  |  | Female | 0 | 1 |
|  | *Trem2* * *R47H* <sup>HSS</sup> Het<br>vs. B6 | Male | 29 | 11 |
|  |  | Female | 2 | 1 |
|  | *Trem2* * *R47H*<br>vs. B6 | Male | 3 | 0 |
|  |  | Female | 0 | 0 |
|  | *Trem2* * *R47H* Het<br>vs. B6 | Male | 9 | 21 |
|  |  | Female | 0 | 2 |
|  | *Trem2* * *R47H*<br>vs. *Trem2* * *R47H* <sup>HSS</sup> | Male | 3 | 1 |
|  |  | Female | 3 | 2 |

**Figure 3.**
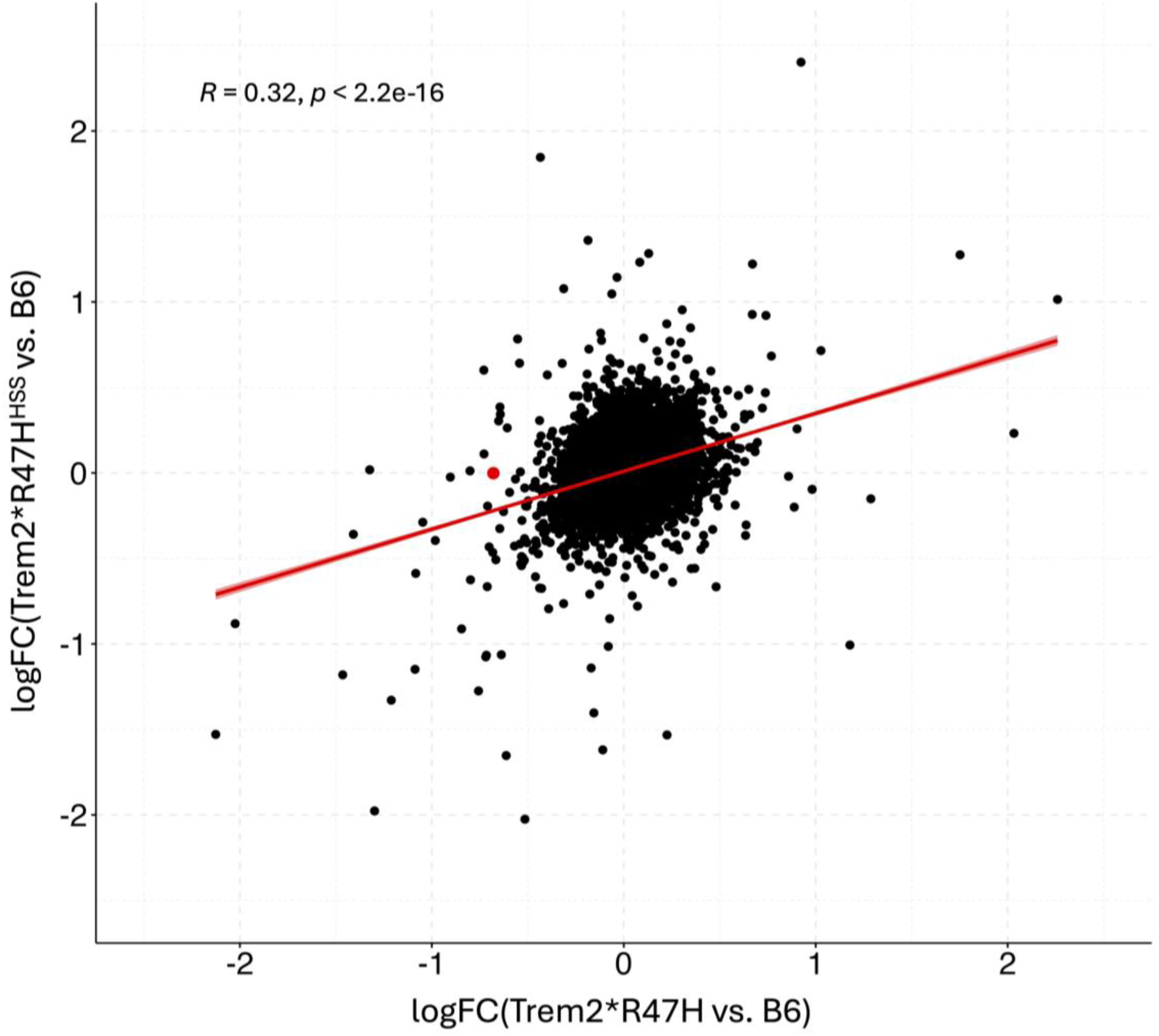
Trem2*R47H and Trem2*R47H^HSS^ mice have correlated gene expression. All transcripts from Trem2*R47H and Trem2*R47H^HSS^ mice were plotted using a scatter plot and correlation was calculated for sex-corrected residual values from 12-month-old Trem2*R47H and Trem2*R47H^HSS^ mice (p < 2.2×10^-16^). The only gene with significant fold change (adj pvalue < 0.05) was Trem2 and is shown as a red dot. The red line shows the slope of the correlation and R and p values for the correlation are shown in the top left-hand corner of the graph.

### 3.3 Trem2*R47H and Trem2*R47H^HSS^ mice display similar correlations with human late onset Alzheimer’s disease transcriptional profiles

Given the interest in Trem2*R47H and Trem2*R47H^HSS^ as potential models for LOAD, we wanted to determine how these mice compared to expression modules previously identified in LOAD patients. To achieve this, we determined the correlation across the genes expressed both in our mouse models and in human cohorts for each of the thirty predefined co-expression modules (see Methods) [31]. Results from a sex-controlled analysis of 12-month-old mice indicated that the Trem2*R47H model was correlated primarily with disease signatures in Consensus Cluster A (ECM organization), Consensus Cluster C (neuronal system), and Consensus Cluster D (cell cycle, NMD) **(Figure 4A)**. There were also some correlations with specific modules in Consensus Cluster B (immune system) and Consensus Cluster E (organelle biogenesis, cellular stress response). The Trem2*R47H^HSS^ model correlated in an overlapping manner except for a lack of correlation in all the Consensus Cluster A and E modules and some of the consensus Cluster C and D modules. A direct comparison between the Trem2*R47H and Trem2*R47H^HSS^ models confirmed that many of these modules were directly influenced by R47H genotype, most notably those associated with Consensus cluster A (ECM organization). Also, direct comparison showed increased positive association of Trem2*R47H mice with Consensus Cluster D which is known to contain signatures associated with oligodendrocytes and myelination. Of note, both models had anti-correlation associated with the STGblue module of Consensus Cluster B (Immune system). Analysis stratified by sex indicated that the Trem2*R47H male mice similarly are overall slightly more correlated with LOAD signatures than Trem2*R47H^HSS^ male mice, however, female mice displayed the opposite pattern and were better correlated in the Trem2*R47H^HSS^ female mice (Supplemental Figure 5). Additionally, four-month-old mice showed less correlation with LOAD signatures overall and *Trem2* heterozygous mice in some modules

**Figure 4.**
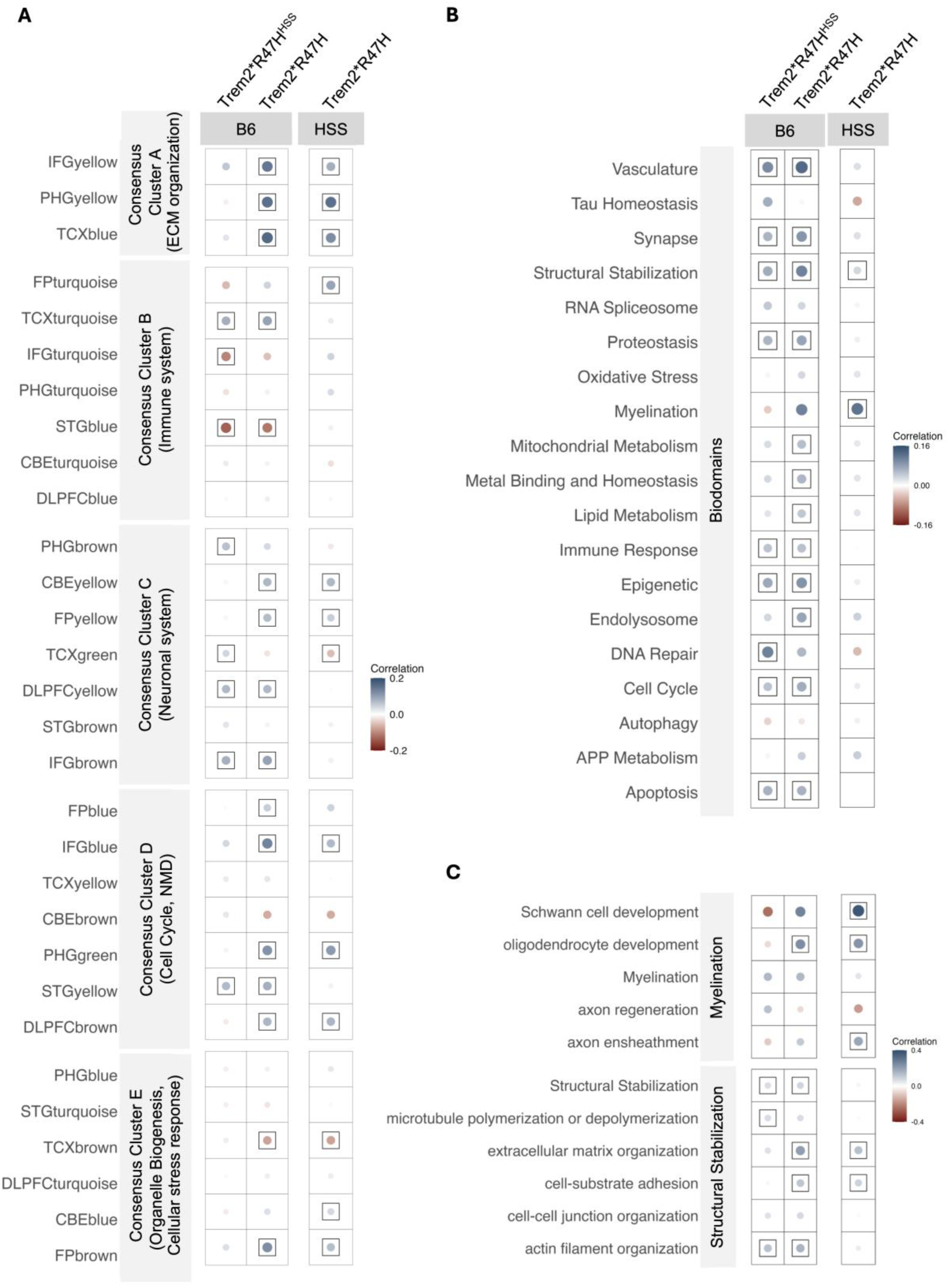
Alzheimer’s-related gene expression modules and biological pathways identified in both Trem2*R47H mice with cryptic splice site and the humanized splice sight are similar, but some can be attributed to loss of expression. A) Dot plot of trans-species correlation analysis comparing gene expression modules enriched in brains from human AD patients to the differential gene expression of mouse models controlled for sex using a linear regression model. Each dot represents a module where the size of the dot indicates the correlation value, the color indicates correlation (blue = correlated, red = anti-correlated), and a black box around the dot indicates statistical significance (p < 0.05). B) Dot plot of trans-species correlation analysis comparing groups of biological pathways (biodomains) enriched in brains from human AD patients to the differential gene expression of mouse models controlled for sex using a linear regression model. Each dot represents a module where the size of the dot indicates the correlation value, the color indicates correlation (blue = correlated, red = anti-correlated), and a black box around the dot indicates statistical significance (p < 0.05). C) Dotplot of significant results from subdomain analysis. HSS = Trem2*R47H^HSS^ mouse model

### 3.4 Trem2*R47H and Trem2*R47H^HSS^ mice are correlated with inflammatory subtypes of Alzheimer’s disease

While we were able to identify correlation with large gene modules common across all LOAD patients, it is known that LOAD is highly heterogenous both clinically and molecularly. To better understand what LOAD subtypes our *Trem2* mouse models represent we assessed correlation of the mouse models with molecular subtypes of AD previously identified in human cohorts [32, 33]. When sex is regressed out, we noted that R47H mouse correlate well with the ROSMAP A, Mayo A, Mayo B, and MSBB A subtypes reported in the *Milind et al* publication while displaying anti-correlation with all other subtypes (Supplemental figure 6A). The Trem2*R47H^HSS^ model was very similar with the exception that the correlations were less strong and the anti-correlation with MSBB B molecular subtype was not significant. Similar patterns of correlation were noted in subtypes from the *Neff et al.* publication with both *Trem2* mouse models correlating with subtype C1 and C2 (Supplemental figure 6B). When stratified by age and sex the correlation patterns are observed to be strongest in male mice and the pattern is reversed at four months of age (Supplemental figure 6C-D).

### 3.5 Additional biological pathways identified in Trem2*R47H mice associated with structural genes

While we were able to identify the correlation of *Trem2* mouse models with human AD-relevant gene modules, we wanted to better understand which specific biological pathways were being altered. To do so we leveraged the strategic sorting of gene ontology terms into distinct AD-relevant biological categories, known as AD biodomains [34]. Results for Trem2*R47H mice showed correlation with multiple pathways including those related to apoptosis, cell cycle, endocytosis, endolysosome, epigenetics, lipid metabolism, metal binding and homeostasis, mitochondrial metabolism, proteostasis, structural stabilization, synapse, and vasculature **(Figure 4B)**. In comparison, the Trem2*R47H^HSS^ mice showed additional correlation with DNA repair, but lacked correlation with the endolysosome, lipid metabolism, and metal binding and homeostasis. Notably, the direct comparison between Trem2*R47H mice and Trem2*R47H^HSS^ mice was shown to primarily influence the strength of the association with the structural stabilization pathways. Using sub-biodomains we were able to determine the specific altered pathways to be those associated with extracellular matrix (ECM) processes associated with cell-substrate adhesion as well as extracellular matrix organization (**Figure 4C**, Supplemental figure 7). We additionally noted some alterations in subdomains associated with myelination – specifically oligodendrocyte development (**Figure 4C**, Supplemental figure 7). However, we did not note any Trem2*R47H vs. Trem2*R47H^HSS^ correlations associated with the immune sub-biodomains (Supplemental figure 7).

## 4 Discussion

Recent advances in mouse models have aided our understanding of the mechanistic drivers underlying LOAD risk associated with *TREM2*. However, there are still many understudied questions, including the impact of different *TREM2* variants (missense, eQTLs, and splicing variants) on LOAD molecular signatures and pathology. Our goal was to molecularly compare a Trem2*R47H model (known to contain cryptic splicing and partial loss of *Trem2* expression) to the Trem2*R47H^HSS^ mouse model (harboring the R47H missense mutation but no reduction in *Trem2* expression). We hypothesized that both *Trem2* models would demonstrate molecular associations with LOAD signatures and provide suitable models for understanding LOAD pathological mechanisms.

Initial transcriptional analysis suggests altered *Trem2* expression/splicing and *Trem2* loss of function mutation R47H have a similar influence on global gene expression patterns. Differential expression analysis of all transcripts from bulk RNA-seq of brain tissue demonstrated the greatest number of significantly differentially expressed genes were present in comparisons of the *Trem2* models to B6 mice. In contrast, there were very few significantly differentially expressed transcripts when directly comparing Trem2*R47H and Trem2*R47H^HSS^ mice and there was a significant correlation between *Trem2* models when comparing all transcripts (unfiltered for significance). While *Trem2* itself did not reach statistical significance in sex-corrected linear regression analysis of Trem2*R47H vs. Trem2*R47H^HSS^ gene expression, the fold-change was in the expected direction and magnitude. Additionally, differential expression analysis of combined males and females resulted in only one significant gene, *Trem2*, identified in comparison of Trem2*R47H vs. B6. As such, we concluded that the significance threshold for adjusted p-value was not reached during multiple comparisons due to high variability in *Trem2* expression (likely exacerbated by the low overall expression of this gene under physiological conditions in the brain). The few genes that were differentially expressed in Trem2*R47H mice as compared to Trem2*R47H^HSS^ mice were associated with extracellular matrix (Col3a1) and metabolism (Hmgcs1, Ldlr, mt-Nd4l). While both *Trem2* loss of function models do not share many overlapping DEGs, the overall trend in expression is highly correlated, especially when considering certain subsets of genes. The human AMP-AD modules that were correlated in both Trem2*R47H and Trem2*R47H^HSS^ mice include genes associated with the immune system, neuronal system, and cell cycle. In addition, both *Trem2* models showed similarity in correlation with LOAD molecular subtypes demonstrating correlation with molecular subtypes identified previously by *Milind et al.* and *Neff et al.* as inflammatory due to strong immune cell signatures [32, 33].

Notably, alternative splicing and partial loss of *Trem2* expression in the Trem2*R47H model appears to result in a slightly stronger LOAD-associated molecular phenotype. While both models showed the similar correlation patterns with AMP-AD modules and AD molecular subtypes, we noted that the Trem2*R47H model overall had greater correlation coefficient values and number of significantly correlated modules than the Trem2*R47H^HSS^ model, suggesting that there may be increased LOAD signature associated with reduced *Trem2*expression. Direct comparison between Trem2*R47H and Trem2*R47H^HSS^ transcriptional signatures in the biodomain analysis suggests that this increased pathogenic profile is driven primarily by additional impacts on ECM and myelination pathways. More broadly, these results were confirmatory of the positive correlation identified in AMP-AD Consensus Cluster A (containing ECM genes) and Consensus Cluster D (containing oligodendrocyte/myelination genes). Specific sub-biodomains identified as significantly correlated in the Trem2*R47H model were the structural biology domains associated with cell substrate adhesion and extracellular matrix organization and one of the myelination sub-biodomains (oligodendrocyte development). The two structural biology sub-biodomains contain several genes previously identified in a proteomic matrisome signature module (module 42) from human LOAD brains [35]. The overlap is significant for both the cell substrate adhesion (p = 0.00014) and extracellular matrix organization (p = 6×10^-13^) sub-biodomains as determined by Fisher’s exact test and includes AD risk genes such as *App*, *Apoe*, *Mdk*, and *Smoc1*.

These findings are aligned with previous *Trem2* studies and highlight potential pathogenic mechanisms of *TREM2*. Several studies of *Trem2* KO mice have also identified significant alterations to ECM molecular signatures and myelination-related processes [15, 16, 36]. Mouse studies have confirmed that *Trem2* is integral in regulating both the clearance of damaged myelin as well as activation of oligodendrocytes to encourage remyelination [9, 16, 36]. *TREM2* is also associated with differentiation of microglial subtypes, and lack of *TREM2* signaling can lead to an decrease in damage associated microglia which may play an important role in clearance of amyloid plaques within the brain [17, 18, 36, 37]. Thus, it is possible that the ECM-related signatures may be generated in part due to synaptic and axonal damage in neurons related to aberrant synaptic pruning and myelination processes. It is also important to note that while Trem2*R47H mice appear to have slightly increased correlation with LOAD-signatures overall, there are also some modules/biodomains with less correlation or anti-correlation. For example, while the Trem2*R47H^HSS^ model is correlated with the DNA repair biodomain, Trem2*R47H is not – this may represent a Trem2*R47H effect that was masked by low *Trem2* expression. These seemingly contradictory findings can highlight another important reality – the nuanced role of *TREM2* in contributing to both AD risk and protection. Prior studies in mice have identified *Trem2* loss of expression as both pathogenic and protective depending on the background of the model [17, 38].

While we believe these findings are significant, there are some limitations to this work. Most notably, we are comparing *Trem2* models in the absence of any neuropathology (amyloid/tau) or inflammation. Published work by the MODEL-AD center at the University of California Irvine demonstrates that Trem2^R47H^ normal splice-site (NSS) mice crossed with 5xFAD mice provide additional insight into the effect of Trem2 loss of function on inflammation response in neurons and glial cells under neurodegenerative conditions [39]. Thus, it is likely that crossing the Trem2 models presented here with other mouse models for AD could provide us with additional insights into Trem2-mediated pathology. It is also possible that LOAD2 mice, which contain both the Trem2*R47H and humanized APOE, may provide a useful resource for placing Trem2 in the context of AD pathology given known interaction between these genes [40–42]. While Trem2*R47H^HSS^ mice were humanized at the cryptic splice site, none of the mouse models analyzed to date contain a fully humanized Trem2 locus. For greater relevance to human biology, it would be useful to perform similar analysis on newer models of *Trem2*\*R47H containing fully humanized *Trem2* loci. A newer series of gene replacement models contains humanization of *Trem2* and Trem-like proteins around the Trem2 locus which have also been implicated in AD risk, including both a wild-type *TREM2* model (JAX #39172) and a version with the R47H variant (JAX #39173). Overall, it will be important to continue similar analysis on a variety of Trem2 mouse models to place these findings in the broader context of available models [43].

In conclusion, we have demonstrated that both Trem2*R47H and Trem2*R47H^HSS^ mouse models represent viable models for studying the influence of *TREM2* in LOAD pathogenic mechanisms. Our findings suggest that the R47H missense mutation and *Trem2* loss of expression have a similar impact on LOAD-associated signatures and molecular pathways with a few specific differences largely impacting ECM and myelination processes. While the Trem2*R47H^HSS^ model provides scientists with an additional model suited for understanding the unique impacts of the R47H missense mutation, Trem2*R47H mice remain useful for modeling analogous mutations in the human population that lead to altered *Trem2* expression and splicing. As such these mouse models warrant further study to contribute to our overall understanding of *TREM2* in AD pathogenesis.

## Abbreviations

AD: Alzheimer’s disease
AMP-AD: Accelerating Medicines Partnership Program for Alzheimer’s Disease
ECM: extracellular matrix
HSS: humanized splice site
IU/JAX/PITT: Indiana University, The Jackson Laboratory, and the University of Pittsburgh
LOAD: late onset Alzheimer’s disease
MODEL-AD: Model Organism Development and Evaluation for Late-onset Alzheimer’s Disease
NSS: normal splice-site
TPM: transcripts per million
TREM2: Triggering receptor expressed on myeloid cells 2

## 5 Acknowledgements

We gratefully acknowledge the contribution of the Genetic Engineering Technologies Service at The Jackson Laboratory for expert assistance with the work described in this publication.

## 8 Ethics statement

The animal study was reviewed and approved by the Jackson Laboratory Animal Use Committee (IACUC-18051-2).

## 9 Data Availability Statement

Bulk RNA-seq data have been deposited at the AD Knowledge Portal (synID: TBD) and are publicly available as of the date of publication. All other data generated during this study are included in this published article and its supplementary information files.

