## Supplemental figures for "Trem2^R47H^ and reduced *TREM2* expression both mimic human Alzheimer’s disease signatures in mice"

### Supplementary Figure 1

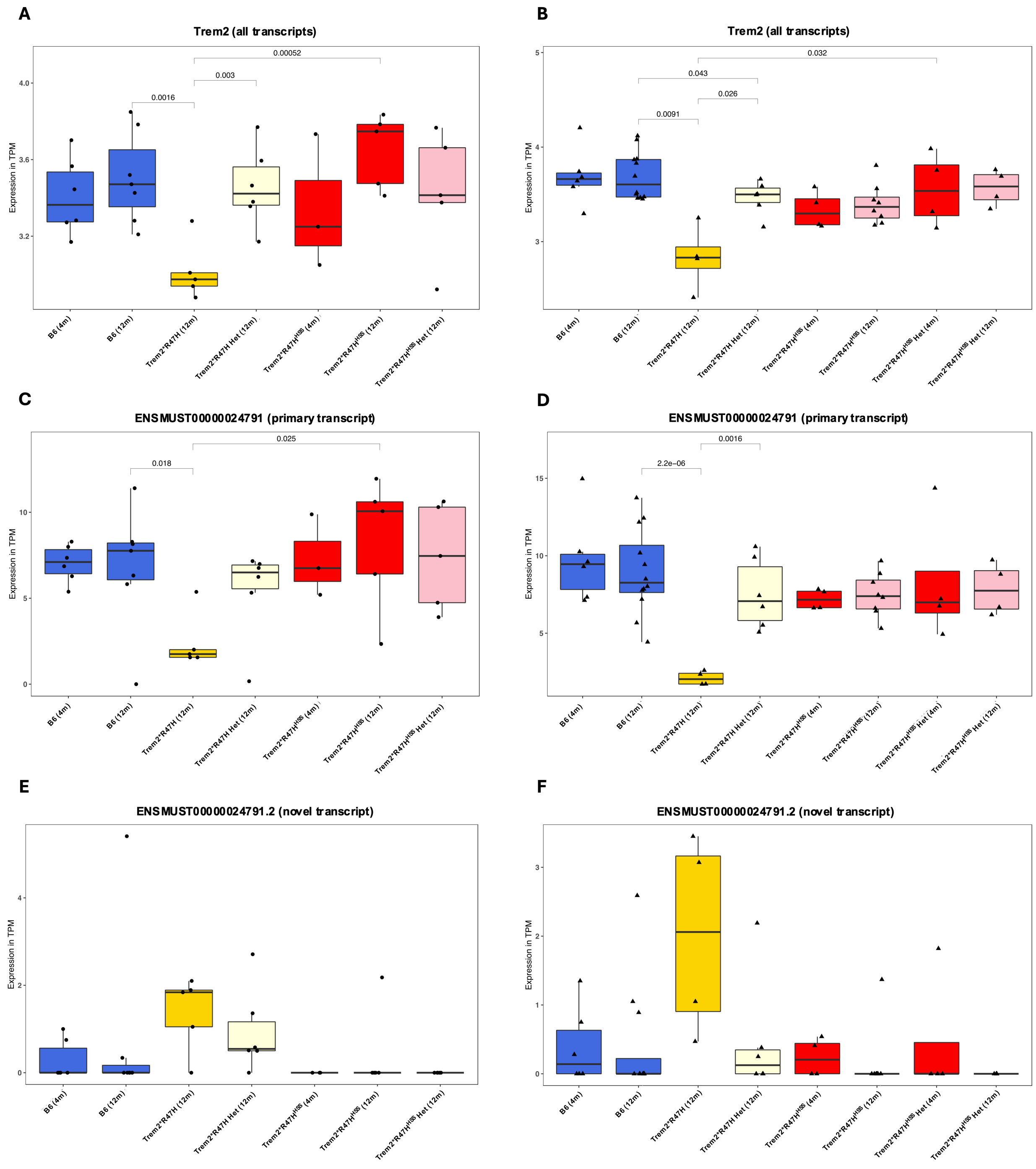

**Supplementary Figure 1. Trem2 isoform expression across mouse age, sex, and genotype.** Expression levels of all Trem2 transcripts (A, B), Trem2 primary transcript (C,D), and Trem2 novel truncated transcript (E,F) stratified by male (A,C,E) and female (B,D,F) mice as well as age (4m, 12m) and genotype (Het = heterozygous). Significant p-values represented by comparison bar and exact p-value (all other comparisons insignificant for $p < 0.05$).

### Supplementary Figure 2

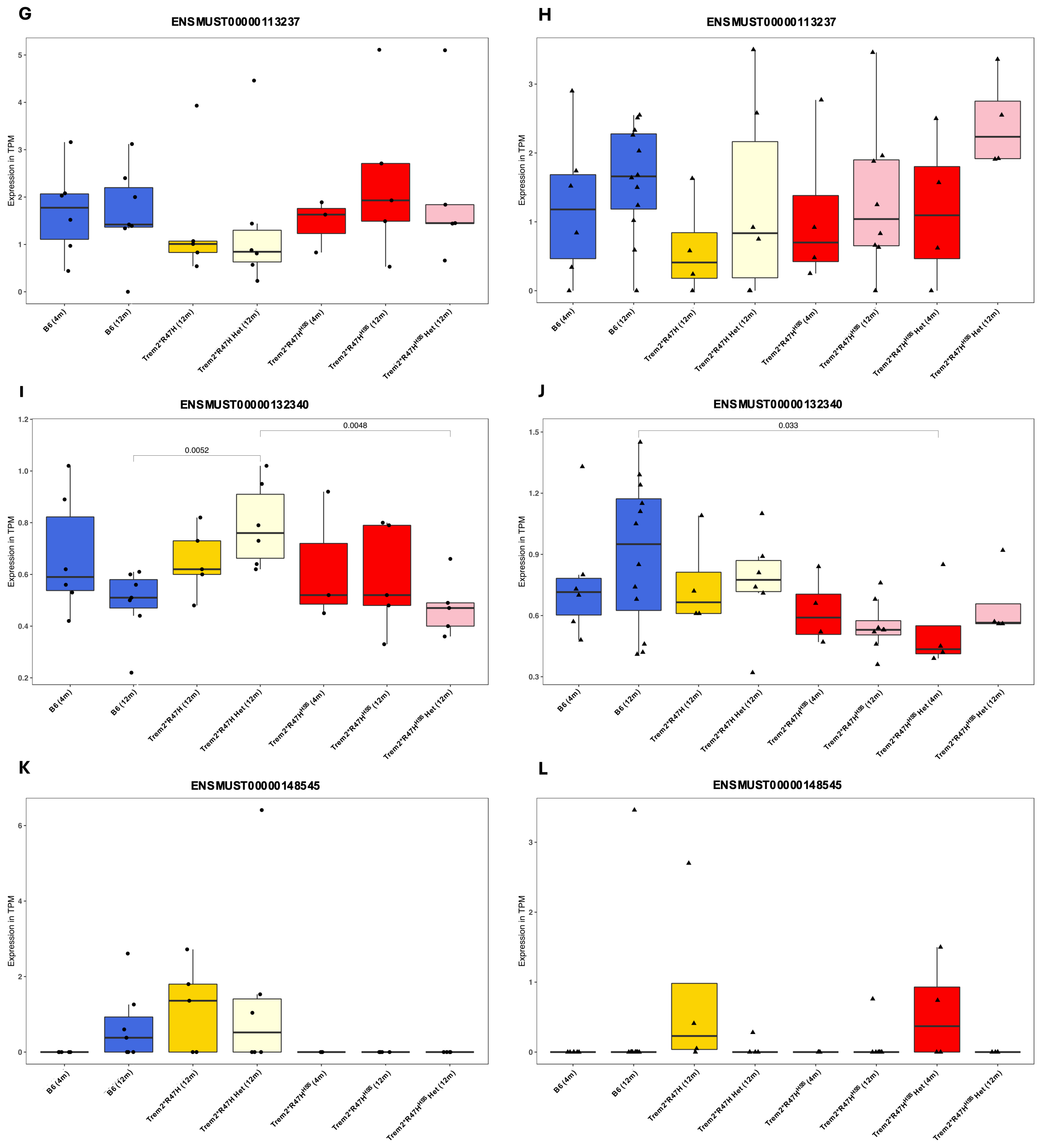

**Supplementary Figure 2. Trem2 isoform expression across mouse age, sex, and genotype (cont.).** Expression levels of Trem2 transcript ENSMUST00000113237 (A, B), ENSMUST00000132340 (C,D), and ENSMUST00000148545 (E,F) stratified by male mice (A,C,E), female mice (B,D,F), age (4m, 12m), and genotype (Het = heterozygous). Significant p-values represented by comparison bar and exact p-value (all other comparisons insignificant for $p < 0.05$).

#### Supplementary Figure 3

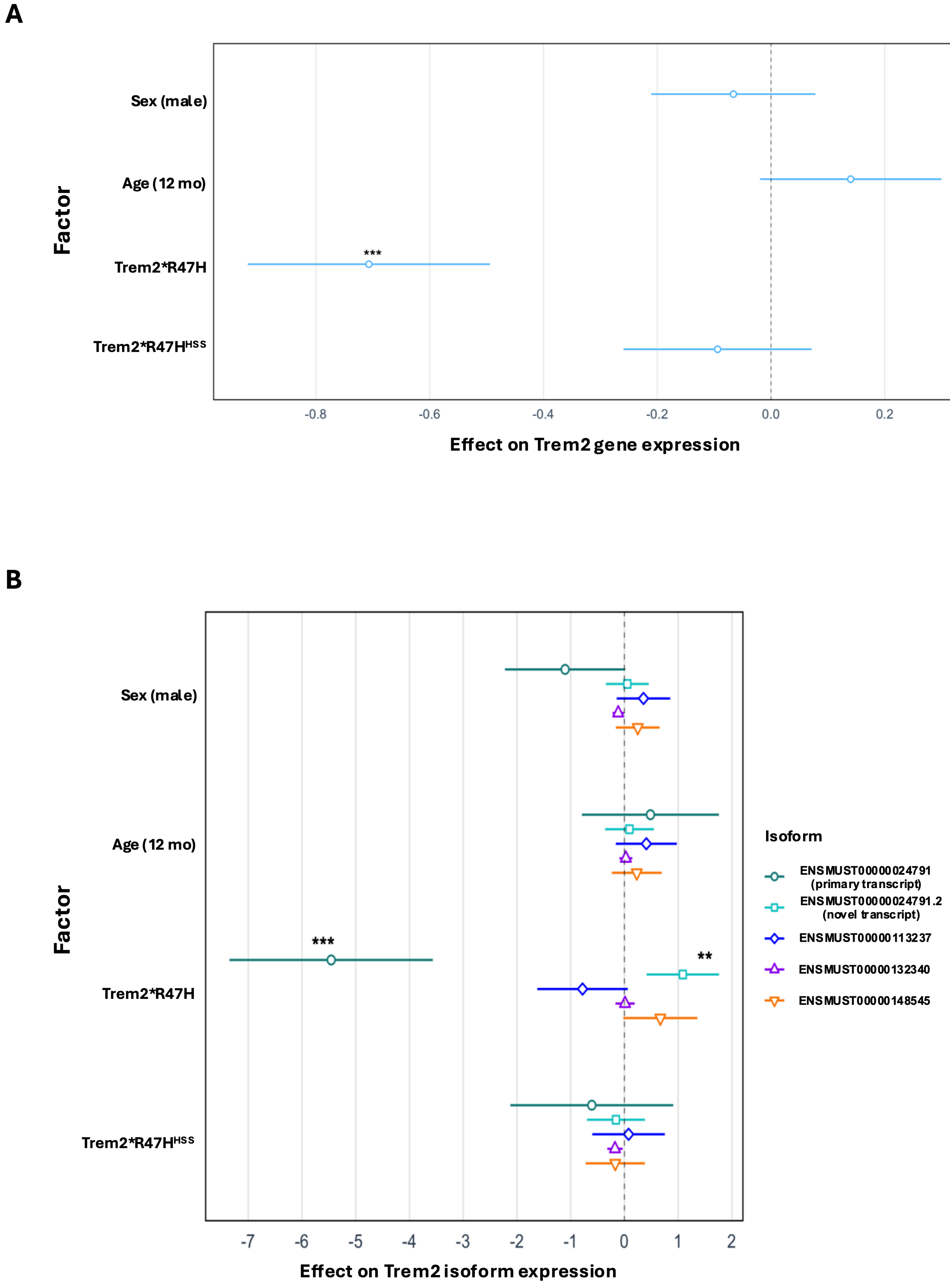

**Supplementary Figure 3. Effect analysis of Trem2 expression.** A) Effect analysis of sex, age, Trem2*R47H genotype, and Trem2*R47H$^{HSS}$ genotype on expression of all Trem2 transcripts 2) Effect analysis of sex, age, Trem2*R47H genotype, and Trem2*R47H$^{HSS}$ genotype on expression of individual Trem2 transcripts.

**Supplemental Figure 4**

**A**

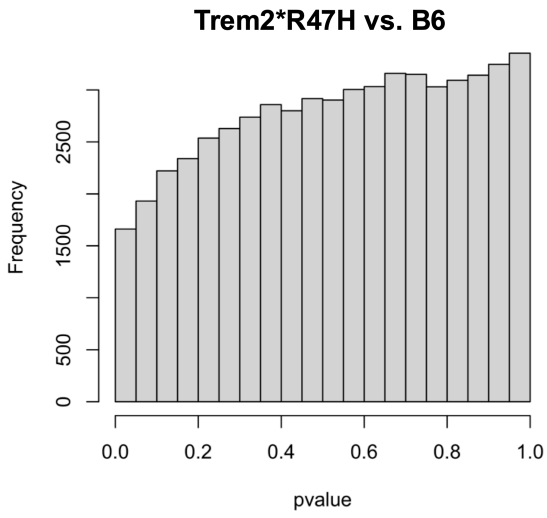


**B**

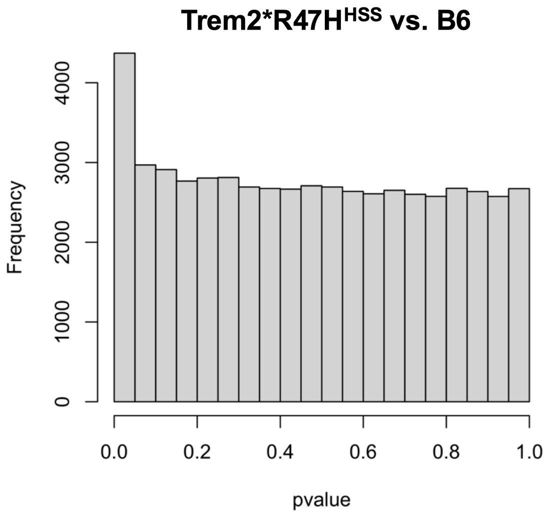


**C**

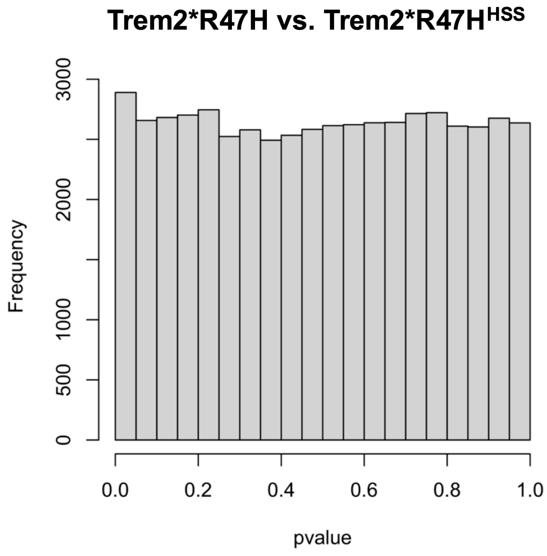

**Supplemental Figure 4. Raw p-value distribution of transcripts from differential expression analysis.** Raw p-values from stratified male/female analysis combined prior to graphing.

### Supplemental Figure 5

A

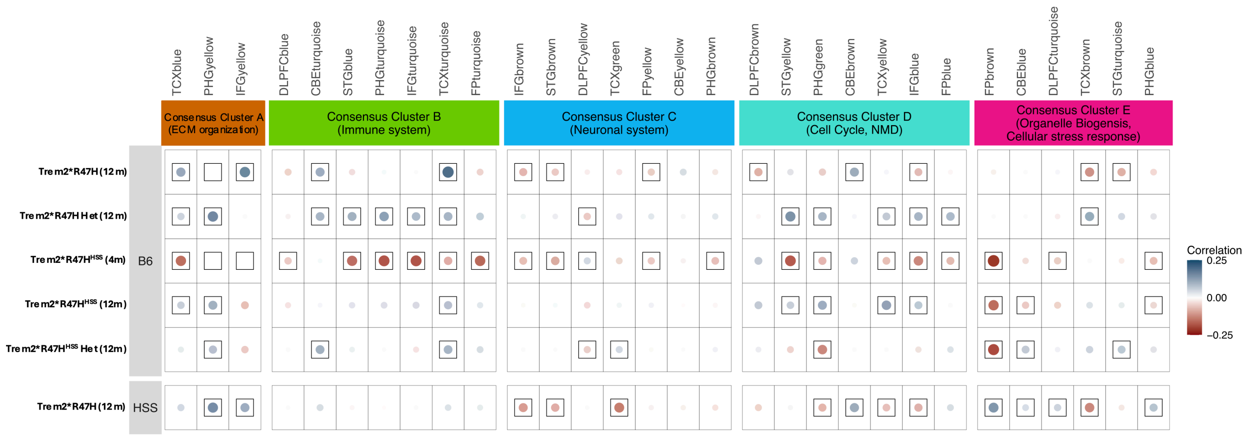


B

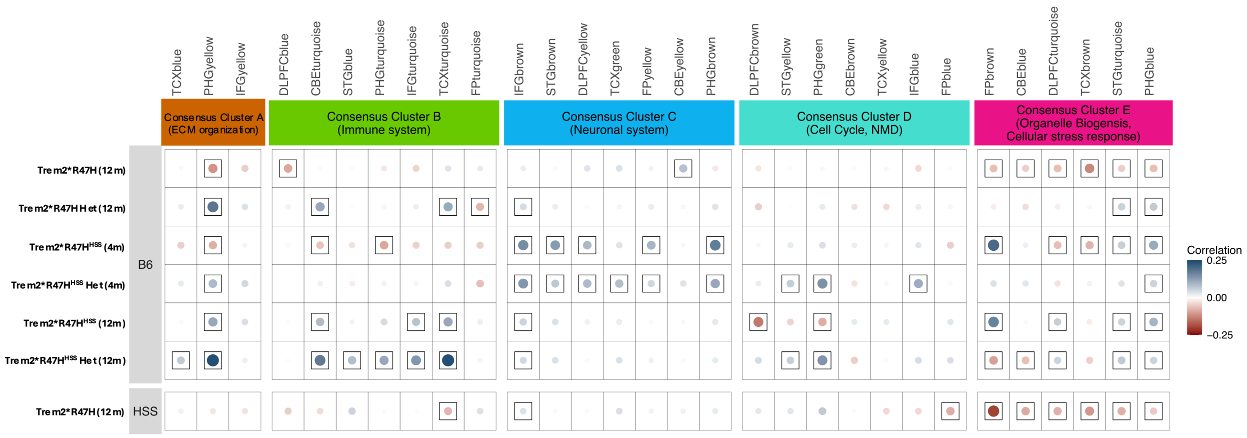

**Supplemental Figure 5. Human AMP-AD module correlation with Trem2 mouse models across age, sex, and genotype.** Trans-species correlation analysis comparing gene expression modules enriched in brains from human AD patients to the differential gene expression of mouse models stratified by A) male and B) female mice. Results plotted as dot plot where each dot represents a module where the size of the dot indicates the correlation value, the color indicates correlation (blue = correlated, red = anti-correlated), and a black box around the dot indicates statistical significance ($p < 0.05$). HSS = Trem2*R47H$^{HSS}$ mouse model

**Supplemental figure 6**

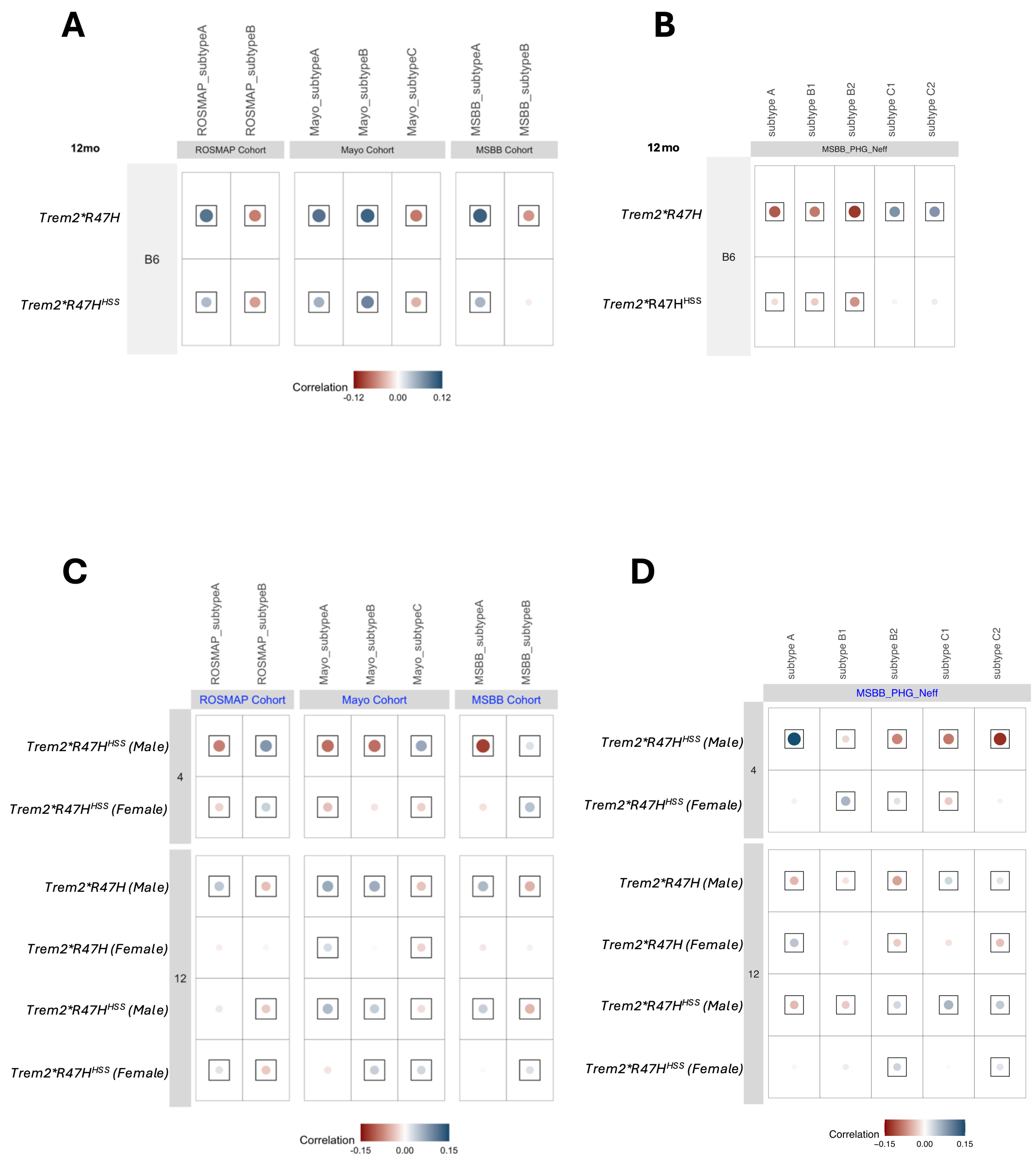

**Supplemental figure 6. Human AD subtype correlation with Trem2 mouse models across age, sex, and genotype.** Trans-species correlation analysis of gene expression enriched in brains of a subset of AD patients (vs. healthy controls) compared to the differential gene expression of Trem2*R47H and Trem2*R47H$^{HSS}$ mice (vs. B6). Differential expression results from linear regression analysis correcting for sex in 12mo old mice were compared to A) *Nikhil et al.* and B) *Neff et al.* subtypes. Results plotted as dot plot where each dot represents a module where the size of the dot indicates the correlation value, the color indicates correlation (blue = correlated, red = anti-correlated), and a black box around the dot indicates statistical significance ($p < 0.05$). Differential expression results from sex-stratified analysis were also compared to C) *Nikhil et al.* and D) *Neff et al.* subtypes.

#### Supplemental figure 7

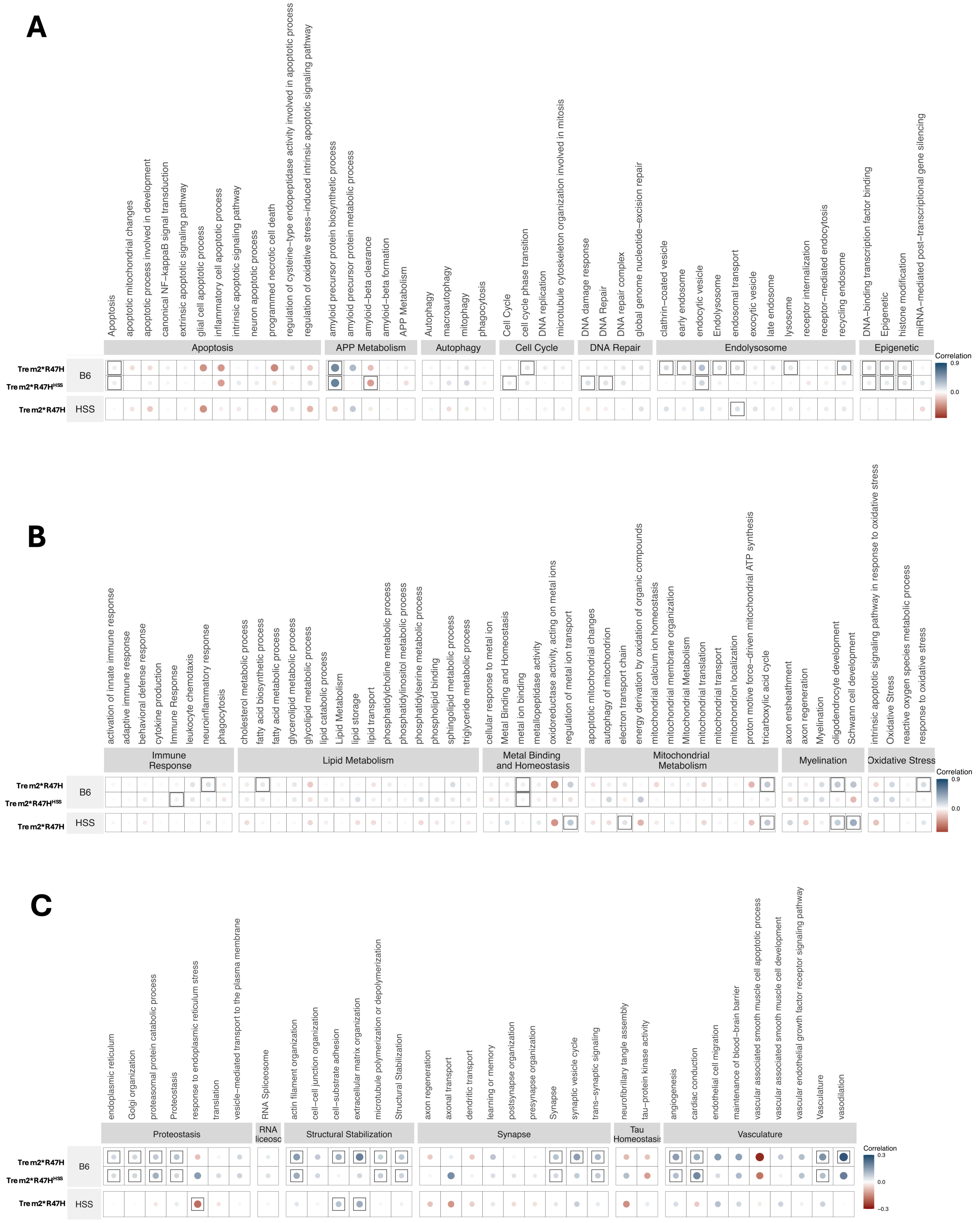

**Supplemental figure 7.** Dot plots of trans-species correlation analysis comparing sub-groups of biological pathways (sub-biodomains) enriched in brains from human AD patients to the differential gene expression of mouse models at 12 months controlled for sex using a linear regression model. Each dot represents a module where the size of the dot indicates the correlation value, the color indicates correlation (blue = correlated, red = anti-correlated), and a black box around the dot indicates statistical significance ($p < 0.05$). Sub-biodomains for the following categories are included: A) apopotosis, APP metabolism, autophagy, cell cycle, DNA repair, endolysosome, epigenetics, B) immune response, lipid metabolism, metal binding and homeostasis, mitochondrial metabolism, myelination, oxidative stress, C) proteostasis, RNA spliceosome, structural stabilization, synapse, tau homeostasis, and vasculature. HSS = Trem2*R47H$^{HSS}$ mouse model
